# Tol-Pal system and Rgs proteins interact to promote unipolar growth and cell division in *Sinorhizobium meliloti*

**DOI:** 10.1101/2020.02.13.948760

**Authors:** Elizaveta Krol, Hamish C. L. Yau, Marcus Lechner, Simon Schäper, Gert Bange, Waldemar Vollmer, Anke Becker

**Author notes:** Simon Schäper, Instituto de Tecnologia Química e Biológica António Xavier, Oeiras, Portugal Hamish C.L. Yau, Newcastle University Faculty of Science, Agriculture and Engineering, NE1 7RX, Newcastle upon Tyne, United Kingdom.

## Abstract

*Sinorhizobium meliloti* is an α-proteobacterium belonging to the Rhizobiales. Bacteria from this order elongate their cell wall at the new cell pole, generated by cell division. Screening for protein interaction partners of the previously characterized polar growth factors RgsP and RgsM, we identified the inner membrane components of the Tol-Pal system (TolQ and TolR) and novel Rgs (rhizobial growth and septation) proteins with unknown functions. TolQ, Pal and all Rgs proteins, except for RgsE, were indispensable for *S. meliloti* cell growth. Six of the Rgs proteins, TolQ and Pal localized to the growing cell pole in the cell elongation phase and to the septum in pre-divisional cells, and three Rgs proteins localized to growing cell pole only. The FtsN-like protein RgsS contains a conserved SPOR domain and is indispensable at the early stages of cell division. The components of the Tol-Pal system were required at the late stages of cell division. RgsE, a homolog of the *Agrobacterium tumefaciens* growth pole ring protein GPR, has an important role in maintaining the normal growth rate and rod cell shape. RgsD is a novel periplasmic protein with the ability to bind peptidoglycan. Analysis of the phylogenetic distribution of novel Rgs proteins showed that they are conserved in Rhizobiales and mostly absent from other α-proteobacterial orders, suggesting a conserved role of these proteins in polar growth.

**IMPORTANCE:** Bacterial cell proliferation involves cell growth and septum formation followed by cell division. For cell growth, bacteria have evolved different complex mechanisms. The most prevalent growth mode of rod shaped bacteria is cell elongation by incorporating new peptidoglycan in a dispersed manner along the sidewall. A small share of rod-shaped bacteria, including the α-proteobacterial Rhizobiales, grow unipolarly. Here, we identified and initially characterized a set of Rgs (rhizobial growth and septation) proteins, which are involved in cell division and unipolar growth of *Sinorhizobium meliloti* and highly conserved in Rhizobiales. Our data expand the knowledge of components of the polarly localized machinery driving cell wall growth and suggest a complex of Rgs proteins with components of the divisome, differing in composition between the polar cell elongation zone and the septum.

## INTRODUCTION

The life cycle of unicellular bacteria includes genome replication and approximate duplication of cell size followed by cell division. Increasing the cell volume relies on cell wall growth, which requires elongation of the peptidoglycan (PG) sacculus. Most of the rod-shaped bacteria elongate by incorporating new PG in a dispersed manner along the sidewall, with the filaments of actin homolog MreB providing a scaffold for the elongasome PG biosynthesis machinery (1, 2). The cell division process relies on constriction of the Z-ring that consists of the tubulin homolog FtsZ, a core component of the dynamic divisome complex (3). The *E. coli* divisome comprises more than twenty different proteins, including SPOR domain protein FtsN and the envelope-spanning Tol-Pal complex (4, 5). The latter consists of the inner membrane protein TolQ, the inner membrane-anchored periplasmic proteins TolR and TolA, the outer membrane-anchored protein Pal and the Pal-associated periplasmic protein TolB (5, 6). Both elongasome and divisome include PG synthases and hydrolases, whose activities are tightly regulated in time and space to maintain cell shape and ensure cell wall integrity (6).

Whereas FtsZ-mediated cell division is conserved among most of the bacterial phyla (7), MreB-dependent cell elongation is less ubiquitous. In Gram-positive Streptomyces, Mycobacteria and Actinobacteria, the polar protein DivIVA fulfills a function in scaffolding PG biosynthesis during cell elongation, which results in bipolar cell growth in rod-shaped species and apical growth at hyphae tips in filamentous species (8–10). The Gram-negative α-proteobacterial order Rhizobiales includes species that lost MreB in the course of evolution (11). Unipolar cell elongation in rod-shaped Rhizobiales, characterized by insertion of new cell wall material at the new cell pole generated by cell division, was reported for several members of this order, such as the plant pathogen *Agrobacterium tumefaciens*, the plant symbiont *Sinorhizobium meliloti* and the animal pathogen *Brucella abortus* (11–13). Despite ample evidence for polar cell growth in Rhizobiales, the scaffolding and regulatory factors governing this process remain largely unknown.

In *A. tumefaciens*, the conserved divisome proteins FtsZ, FtsA and FtsW were suggested to regulate transition from polar growth to cell division (14, 15). Cells depleted for these proteins produce branches originating from the septal site (14). The exceptionally large growth pole ring protein GPR, which localizes to the growing cell pole, was shown to be required for normal polar growth and rod cell morphology in *A. tumefaciens* (16). Its overproduction caused formation of ectopic growth zones and cell branching, thus this protein was proposed to constitute a structural component of an organizing center for PG synthesis during polar growth (16). However, its exact function is yet to be determined.

In *S. meliloti*, 7TMR-DISM-cdGMP phosphodiesterase RgsP (Rgs for rhizobial growth and septation) and putative membrane-anchored periplasmic PG metallopeptidase RgsM were identified as important factors for unipolar cell growth (12). Both are essential proteins and conserved in Rhizobiales (12). These proteins localize to sites of zonal PG synthesis at the growing cell pole and septum. Depletion of RgsP or RgsM results in cell growth inhibition and altered muropeptide composition (12).

Here, we expand the knowledge of components involved in the control of unipolar cell growth and division in Rhizobiales. We present nine further *S. meliloti* proteins with unknown functions that localize to sites of zonal PG synthesis, are required for cell growth, and involved in protein-protein interactions with RgsP and RgsM. We further show that Rgs proteins interact with components of the Tol-Pal system, which is localized to sites of zonal PG synthesis and essential for *S. meliloti* cell division.

## RESULTS

### RgsA is essential and localizes to sites of zonal cell wall growth

We previously identified RgsA (SMc00644) as a potential interaction partner of RgsP and RgsM (12) and recently, a massively parallel transposon insertion sequencing (Tn-seq) study suggested large growth impairment caused by transposon insertions in *rgsA* (17). To test for a functional relation between RgsA, RgsP and RgsM, we attempted to knock out *rgsA* in strain Rm2011 *rgsP-egfp*, carrying replacement of *rgsP* by the gene fusion at the native genomic locus. Since this approach failed, we constructed the RgsA depletion strain Rm2011 *rgsP-egfp rgsA*^dpl^. To enable RgsA depletion, the chromosomal *rgsA* gene was deleted in presence of a plasmid-borne copy of this gene controlled by its native promoter in Rm2011 *rgsP-egfp*. Replication of this single copy plasmid pGCH14 was mediated by a *repABC* operon whose expression was dependent on IPTG (18). Omitting IPTG from the growth medium of this RgsA depletion strain resulted in loss of normal RgsP localization and growth inhibition (Fig. 1A, 1B). In contrast, with IPTG added to the medium, the RgsA depletion strain grew normally (Fig. 1B). RgsA-depleted cells were slightly shorter and wider than the wild type cells and showed increased cell curvature (Fig. 1A, Table 1). This cell morphology was different from the shorter, rounded rods, observed upon RgsP or RgsM depletion (12). Muropeptide analysis of PG isolated from RgsA-depleted cells revealed the following changes in the PG composition: an increase in monomeric tetra- and pentapeptides, and the glycine-containing dimer (TetraTetra(Gly4)), and a decrease in the dimeric (TetraTetra) and trimeric (TetraTetraTetra) cross-linked muropeptides (Fig. 1C, Table 2). These changes in muropeptide composition are similar to those in RgsP- or RgsM-depleted cells reported previously (12). To monitor cellular localization of RgsA by fluorescence microscopy, we replaced *rgsA* with *rgsA-mCherry* at the native genomic location in the *rgsP-egfp* strain. The resulting strain did not differ from the wild type in growth and cell morphology, thus we concluded that the protein fusion was functional. RgsA-mCherry colocalized with RgsP-EGFP at the growing cell pole and the septum (Fig. 1D). RgsA-mCherry and RgsP-EGFP fluorescent foci tracked by time-lapse microscopy showed matching localization dynamics during the cell cycle and during their relocation from the growing pole to the septum (Fig. 1E, Fig. S3), corroborating the suggested close functional relation between the two proteins. These results support the notion that RgsA may constitute a part of a protein complex including RgsP and RgsM, required for *S. meliloti* cell wall growth.

**FIG 1.**
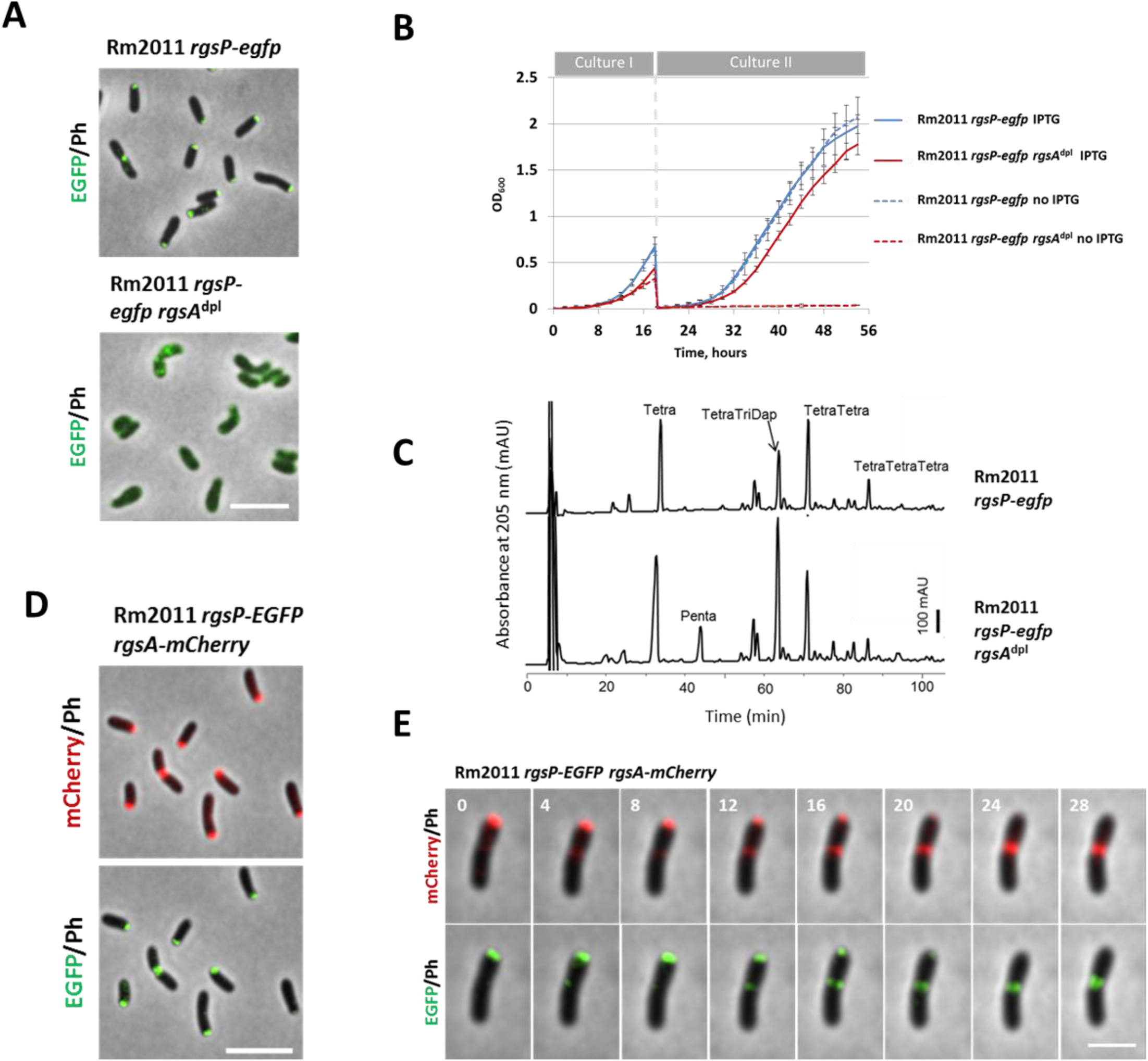
Similar to its interaction partner RgsP, RgsA is essential for growth and normal muropeptide composition, and localizes to the growing cell pole and septum. (A) Fluorescence microscopy images of Rm2011 *rgsP-egfp* and its *rgsA* depletion derivative. Cells from TY cultures, grown for 20 hours without added IPTG are shown. Ph, phase contrast. Bar, 5 µm. (B) Growth of Rm2011 *rgsP-egfp* and its *rgsA* depletion derivative. Cultures I were inoculated at OD_600_ of 0.005 in TY medium either with or without added IPTG, grown for 18 hours and set back to OD_600_ of 0.005 in fresh TY medium with or without added IPTG to start the Cultures II. (C) Muropeptide profiles of PG samples from Rm2011 *rgsP-egfp* and its *rgsA* depletion derivative, grown for 20 hours in TY medium without IPTG. (D) Fluorescence microscopy images of Rm2011 *rgsP-egfp rgsA-mCherry*. Cells from exponentially growing TY cultures are shown. Bar, 5 µm. (E) Time-lapse microscopy images of Rm2011 *rgsP-egfp rgsA-mCherry*. Time is denoted in minutes. Ph, phase contrast. Bar, 2 µm.

**Table 1.** Mean values of cell morphology measurements in cultures, grown without added IPTG for 20-24 hours.

| Strain | Length | Width | Area | Curvature | Roundness | N<sup>1</sup> |
| --- | --- | --- | --- | --- | --- | --- |
| 2011 *rgsP-egfp* | 3.15 | 0.78 | 2.26 | 0.14 | 0.30 | 918 |
| 2011 *rgsP-egfp rgsA<sup>dpl</sup>* | 2.51 | 0.83 | 1.87 | 0.32 | 0.42 | 1000 |
| 2011 *rgsP-egfp rgsB<sup>dpl</sup>* | 2.64 | 0.85 | 2.00 | 0.20 | 0.39 | 1000 |
| 2011 *rgsP-egfp rgsC<sup>dpl</sup>* | 3.00 | 0.88 | 2.44 | 0.18 | 0.36 | 798 |
| 2011 *rgsP-egfp rgsD<sup>dpl</sup>* | 2.16 | 0.95 | 1.77 | 0.21 | 0.50 | 583 |
| 2011 *rgsP-egfp rgsE<sup>dpl</sup>* | 2.27 | 0.93 | 2.12 | 0.77 | 0.55 | 1000 |
| 2011 *rgsP-egfp rgsF<sup>dpl</sup>* | 2.07 | 0.89 | 1.64 | 0.25 | 0.51 | 1000 |
| 2011 *rgsP-egfp rgsG<sup>dpl</sup>* | 2.64 | 0.83 | 2.01 | 0.19 | 0.39 | 813 |
| 2011 *rgsP-egfp rgsH<sup>dpl</sup>* | 2.63 | 0.80 | 1.90 | 0.18 | 0.38 | 1000 |
| 2011 *rgsP-egfp tolQ<sup>dpl</sup>* | 3.64 | 0.77 | 2.63 | 0.19 | 0.29 | 601 |
| 2011 *rgsP-egfp pal<sup>dpl</sup>* | 3.49 | 0.77 | 2.49 | 0.17 | 0.29 | 1000 |

<sup>1</sup> number of analyzed cells.

**Table 2.**
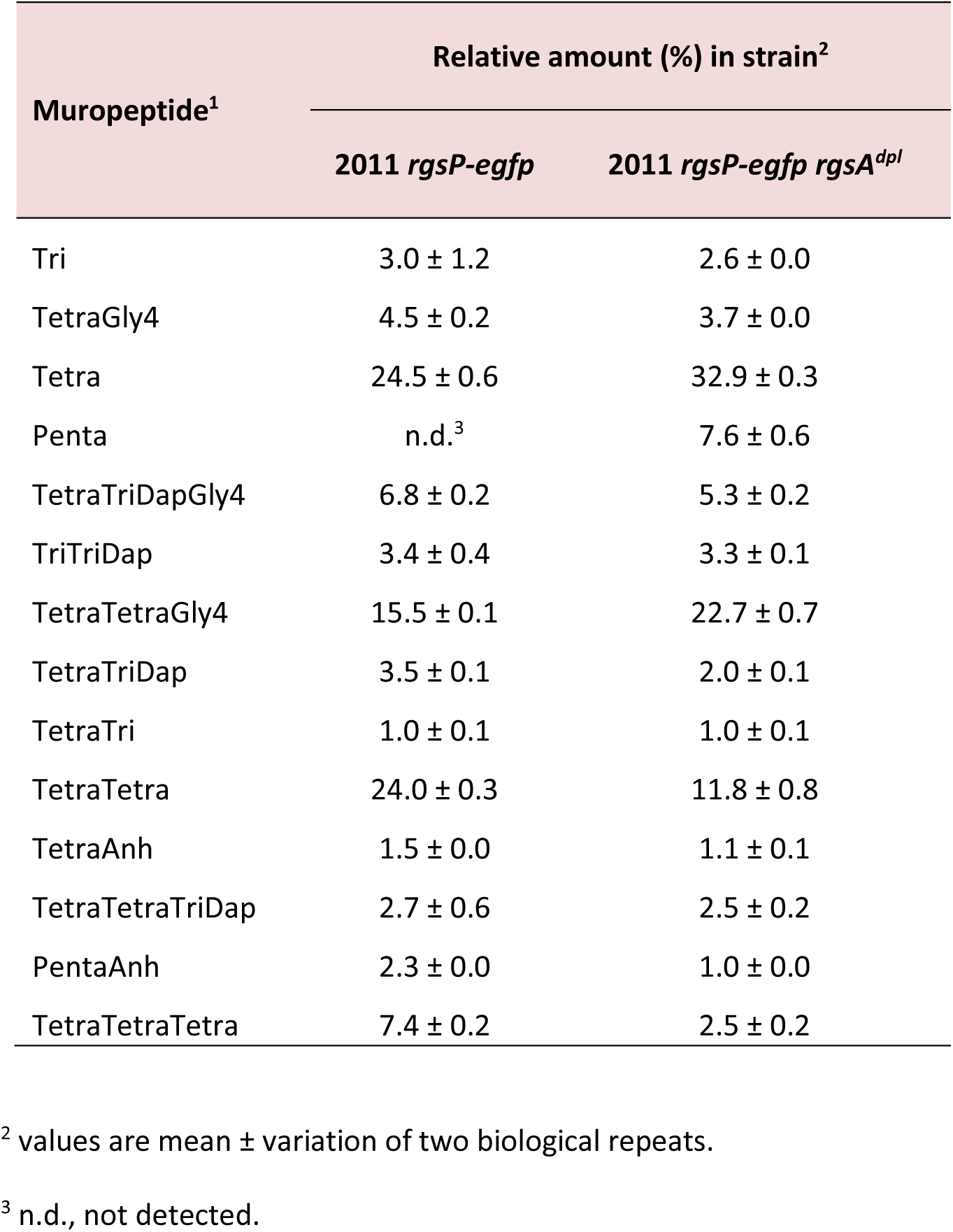
Relative % of muropeptides in PG samples.

### RgsA, RgsM and RgsP, several hypothethical proteins, and components of the Tol-Pal system constitute an interaction network

To test the hypothesis of a functional connection between RgsP, RgsM and RgsA and known cell wall growth factors, we screened for additional protein interaction partners. For this purpose, we replaced the native *rgsA* by a 3xFLAG tagged gene version to generate strain Rm2011 *rgsP-egfp rgsA-3xflag*. This strain was used to immunoprecipitate RgsA-3xFLAG and its putative interaction partners from cells, subjected to cross-linking with formaldehyde. RgsP and RgsM were enriched in the RgsA pulldown sample, implying that these three proteins may form a complex *in vivo* (Fig. 2A, Dataset S1). Moreover, among the enriched proteins we found the TolQ and TolR components of the Tol-Pal system and eight hypothetical proteins encoded by genes of unknown function assigned to the category “essential” or “large growth impairment” in the recent Tn-seq study (17) (Dataset S1). These proteins were named RgsB (SMc04006), RgsC (SMc04010), RgsD (SMc01011), RgsE (SMc00190), RgsF (SMc03995), RgsG (SMc00950), RgsH (SMc00153) and RgsS (SMc02072) (Fig. 2A). RgsE is a homolog of the *A. tumefaciens* growth pole ring protein GPR, recently reported to promote polar growth in this organism (16). Noteworthy, RgsB, RgsD and RgsG were previously identified as putative interaction partners of RgsM, and RgsG as putative interaction partner of RgsP (12). In order to corroborate the identified putative interactions, strains carrying *rgsB-3xflag*, *rgsS-3xflag* and *tolQ-3xflag* were used in pulldown assays. Rgs proteins identified in the RgsB-3xFLAG and RgsA-3xFLAG pulldown samples largely overlapped (Fig. 2A, Dataset S1). Moreover, RgsB was the most strongly enriched co-immunoprecipitated protein in the RgsA-3xFLAG sample and *vice versa*. In the RgsS-3xFLAG pulldown sample, RgsE, TolR and RgsC were among the enriched proteins and pulldown with TolQ-3xFLAG identified TolR, TolA, RgsA, RgsS and RgsE as enriched. Moreover, we used strains generated previously to repeat the pulldown experiments with RgsP-3xFLAG and RgsM-3xFLAG (12). The resulting data corroborated our previous findings and added links between the RgsP-RgsM interaction network and TolQ and RgsS (Fig. 2A, Dataset S1). Collectively, our co-immunoprecipitation results suggest protein-protein interactions between Rgs proteins and the Tol-Pal system.

**FIG 2.**
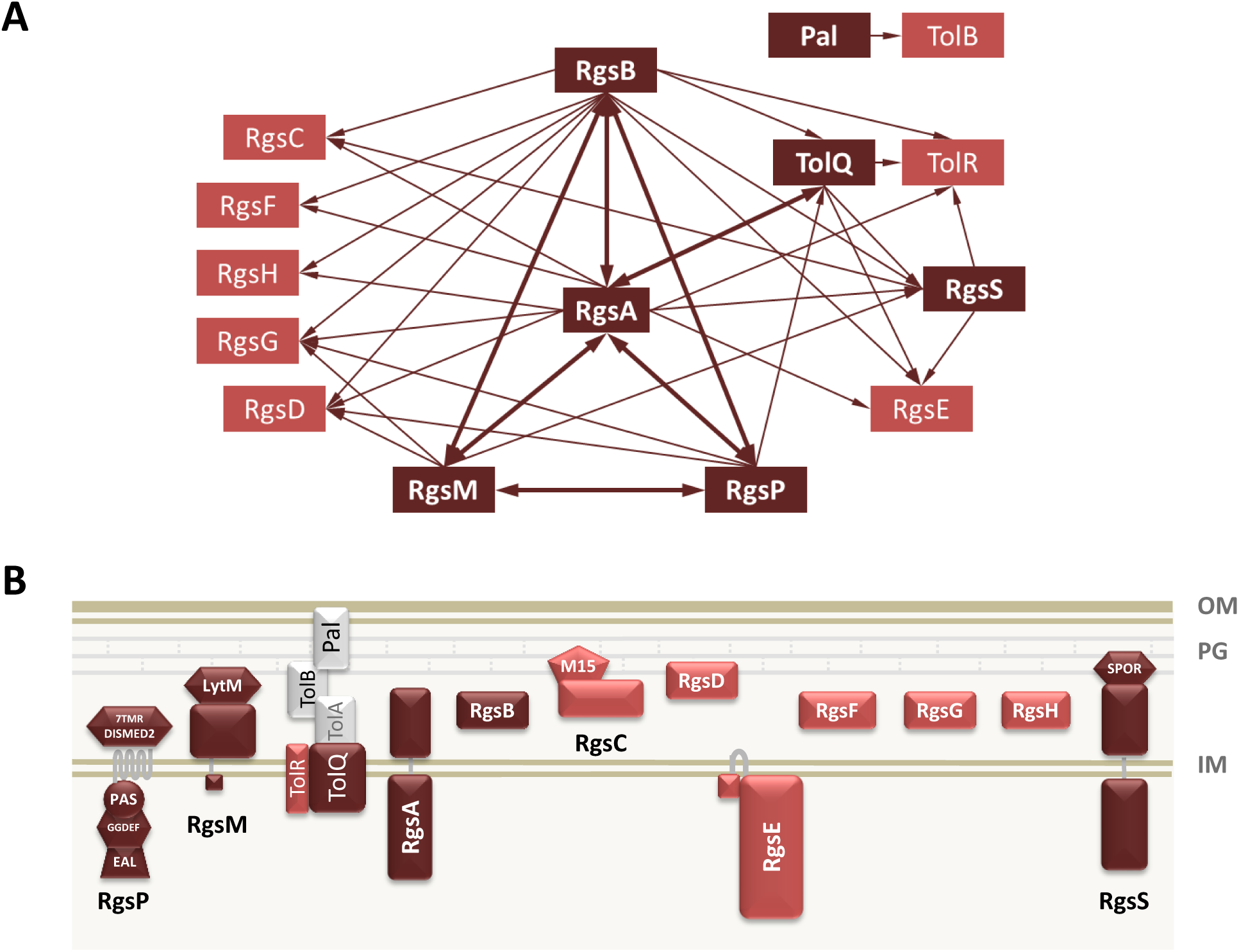
Identification of novel polar growth factors in *S. meliloti*. (A) Summary of putative protein-protein interactions identified in pulldown experiments. Dark red rectangles indicate proteins used as baits. Thick arrows indicate reciprocal pulldowns. (B) Schematic representation of the membrane topology and conserved domains in Rgs proteins and the Tol-Pal system.

We did not find the outer membrane-anchored and periplasmic components of the Tol-Pal system, TolB and Pal, among the interaction partners of TolQ or Rgs proteins. Thus, we carried out a pulldown experiment with a strain carrying *pal-3xflag* at the native genomic location. In this assay, only TolB was strongly enriched, indicating that Pal is unlikely to directly interact with Rgs proteins and inner membrane components of the Tol-Pal system (Fig. 2A, Dataset S1).

### Membrane topology of novel Rgs proteins and their interactions in a bacterial two-hybrid assay

To get first hints on properties of the novel Rgs proteins, we analyzed their protein sequences with the Phobius transmembrane topology prediction tool (19). In RgsA and RgsS, a large N-terminal cytoplasmic and a C-terminal periplasmic domain separated by a transmembrane helix was predicted. RgsE was predicted to be primarily located in the cytoplasm, anchored in the inner membrane via two transmembrane helices (Fig. 2B, Dataset S2). Periplasmic location and signal peptides were predicted for RgsB, RgsD, RgsG, RgsH and RgsF, whereas RgsC was suggested to be an extracytoplasmic protein without a signal peptide (Fig. 2B, Dataset S2).

To validate the predicted membrane topology, we used fusions to the partial *E. coli* alkaline phosphatase PhoA_27-471_, lacking the signal peptide. PhoA is only enzymatically active if located in the periplasm where its activity results in blue staining of the bacterial culture on indicator plates. Rgs protein coding sequences of different lengths were inserted into a plasmid, carrying the PhoA_27-471_ coding sequence, and the resulting plasmids were introduced into *S. meliloti* Rm2011 and *E. coli* S17-1. Periplasmic location, dependent on the signal peptide, was confirmed for RgsB, RgsD, RgsF, RgsG and RgsH, since fusions of the full-length proteins to PhoA_27-47_ produced blue staining of the respective bacterial cultures in both species and fusions of proteins lacking the predicted signal peptide portion did not produce blue staining (Fig. S1A). Furthermore, cytoplasmic location of the N-terminal portion, periplasmic location of the C-terminal portion and location of the transmembrane segment in RgsA and RgsS could be verified, as well as a cytoplasmic location of RgsE and its membrane anchoring (Fig. S1A).

Comparison of the annotated RgsC sequence with homologous sequences in databases suggested a longer protein, with additional 43 amino acids at the N-terminus (Fig. S1B). This increased the length of the protein to 605 amino acids. RgsC_1-605_ fused to PhoA_27-471_ produced blue staining on indicator plates, which suggested periplasmic location of the fusion protein. Thus we used fusions to PhoA to determine the length of the signal peptide. Fusion of the first 35 amino acids of RgsC to PhoA_27-471_ did not result in blue staining in strains bearing the corresponding construct, whereas a similar construct carrying the first 38 amino acids produced the staining (Fig. S1). This indicates that the signal peptide of RgsC is 36 to 38 amino acids long. The cysteine at position 37 might represent an acylation site, typically located directly after the signal peptide cleavage site in lipoproteins (20).

Protein-protein interactions were tested in a bacterial two-hybrid system. We employed a system based on reconstitution of active adenylate cyclase from T18 and T25 fragments upon interaction of the fused proteins of interest. β-galactosidase activity as the signal output produces blue staining on indicator plates with X-Gal. Since adenylate cyclase is a cytoplasmic enzyme, we only fused the T18 and T25 fragments to proteins containing cytoplasmic domains. Strong blue staining indicated interaction of TolQ with RgsM, RgsE and itsef, as well as of RgsA with RgsE and RgsS. Weaker staining was caused by combining RgsM with RgsP, RgsA and itself, RgsA with TolQ and itself, and RgsE with RgsS (Fig. 3). The negative result obtained by combining RgsA with RgsP might be a hint that RgsP-RgsA interaction is indirect, possibly mediated by RgsM. Alternatively, it might be due to unfavorable conformations of RgsA and RgsP proteins fused to the adenylate cyclase fragments.

**FIG 3.**
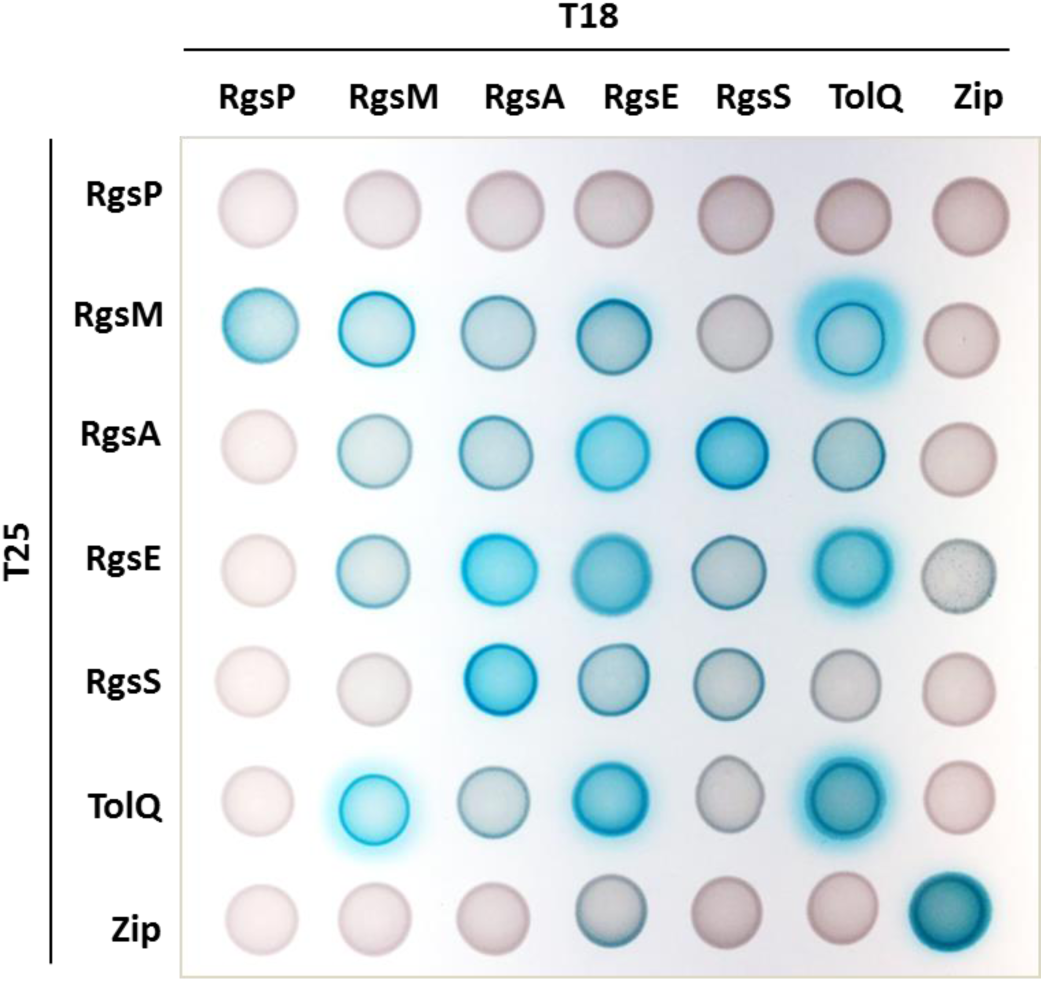
Bacterial two-hybrid analysis. *E. coli* strain BTH101 was co-transformed with the indicated T18 and T25 fusion constructs and 10 µl of the co-transformant cell suspensions were spotted onto LB agar containing X-Gal. Blue staining indicates protein-protein interactions.

### Rgs proteins and Tol-Pal are required for growth of *S. meliloti*

Our attempts to generate knock-out mutants failed for all the novel *rgs* genes, *tolQ*, and *pal*. Thus we constructed depletion strains in the Rm2011 *rgsP-egfp* genetic background. To generate RgsB, RgsC, RgsD, RgsE, RgsF, RgsG, RgsH, RgsS, TolQ, and Pal depletion strains, the encoding gene was either placed under control of an IPTG-inducible promoter or placed under the control of the native promoter on a single-copy plasmid whose replication is IPTG-dependent (see Materials and Methods). Growth of the depletion strains in medium with added IPTG was similar to that of the wild type, whereas without added IPTG growth was inhibited (Fig. 4A). However, RgsE depletion resulted in slow but constant growth, suggesting that RgsE is not essential.

**FIG 4.**
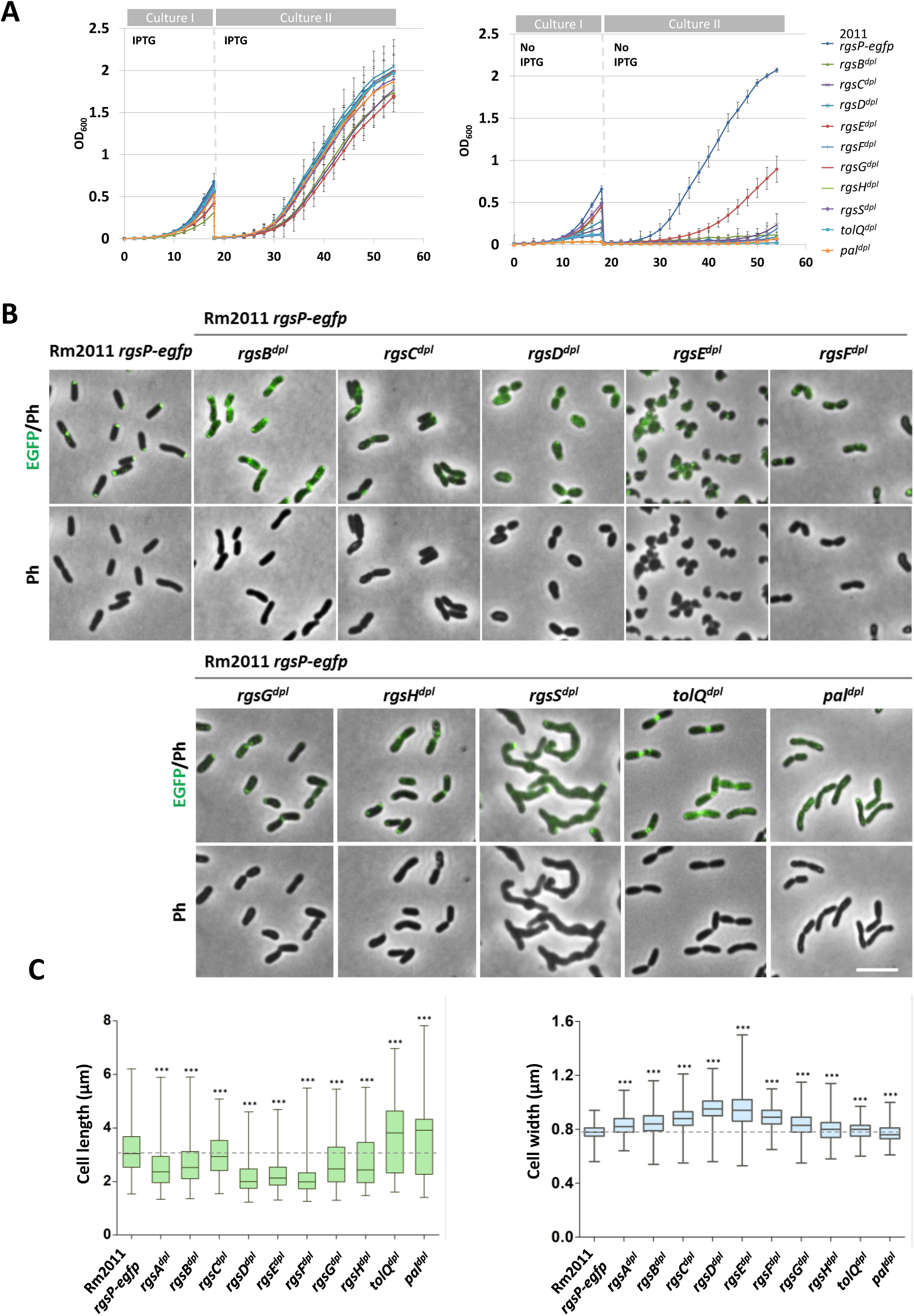
Effects of TolQ, Pal and novel Rgs protein depletion on growth, cell morphology and RgsP localization. (A) Growth of Rm2011 *rgsP-egfp* (Control) and its *rgs* or *tol-pal* depletion derivatives in TY medium. Cultures I either with or without added IPTG were inoculated at OD_600_ of 0.005, grown for 18 hours and used to start Cultures II in fresh TY medium with or without added IPTG at OD_600_ of 0.005. Error bars indicate standard deviation of three biological replicates. (B) Fluorescence microscopy images of Rm2011 *rgsP-egfp*, and its *rgs* or *tol-pal* depletion derivatives, grown in TY without added IPTG for 20-24 hours. Ph, phase contrast. Bar, 5 µm. (C) Box plot diagrams of cell lengths and cell widths. The boxes show 25 % to 75 % percentile, and whiskers show min and max values. Stars indicate statistically significant difference between the mean values calculated for the respective depletion strain and the parental strain Rm2011 *rgsP-egfp* determined by t-test with Welch correction.

In cells, depleted for Rgs or Tol-Pal proteins, the characteristic polar and septal RgsP-EGFP localization was either diminished or lost (Fig. 4B). Moreover, Rgs and Tol-Pal depletion resulted in significant changes in cell length and width, as well as alterations in cell area, roundness and curvature (Fig. 4B, 4C, Table 1). Whereas almost normal rod cell morphology was retained in RgsB, RgsC, RgsG and RgsH depletion conditions (Fig. 4B, 4C), cells depleted for RgsD or RgsF were strongly shortened and rounded, similar to RgsP- and RgsM-depleted cells (12). RgsE-depleted cells were irregularly shaped with strong curvature. These cells were morphologically similar to the *A. tumefaciens* GPR-depleted cells (16). Depletion of RgsS resulted in formation of short, partially branched cell filaments with bulges along the filaments, which may represent swollen septation sites. The filament tips contained RgsP-EGFP fluorescence signal. This implies that cell division but not elongation requires RgsS, since despite unsuccessful cell division, cells were apparently able to elongate further (Fig. 4B). Upon depletion of TolQ or Pal, a strong increase in the mean cell length accompanied by minor changes in cell width was reflective of accumulation of pre-divisional doublets of normally rod-shaped cells, separated by constricted septa (Fig. 4B). This implies that the Tol-Pal system is not crucial for *S. meliloti* cell elongation, despite its localization at the growing cell pole, but is essential for completion of cell division. Electron microscopy of TolQ-depleted cells revealed extended tube-like constricted septa, corroborating a defect in late stages of cell division (Fig. 5).

**FIG 5.**
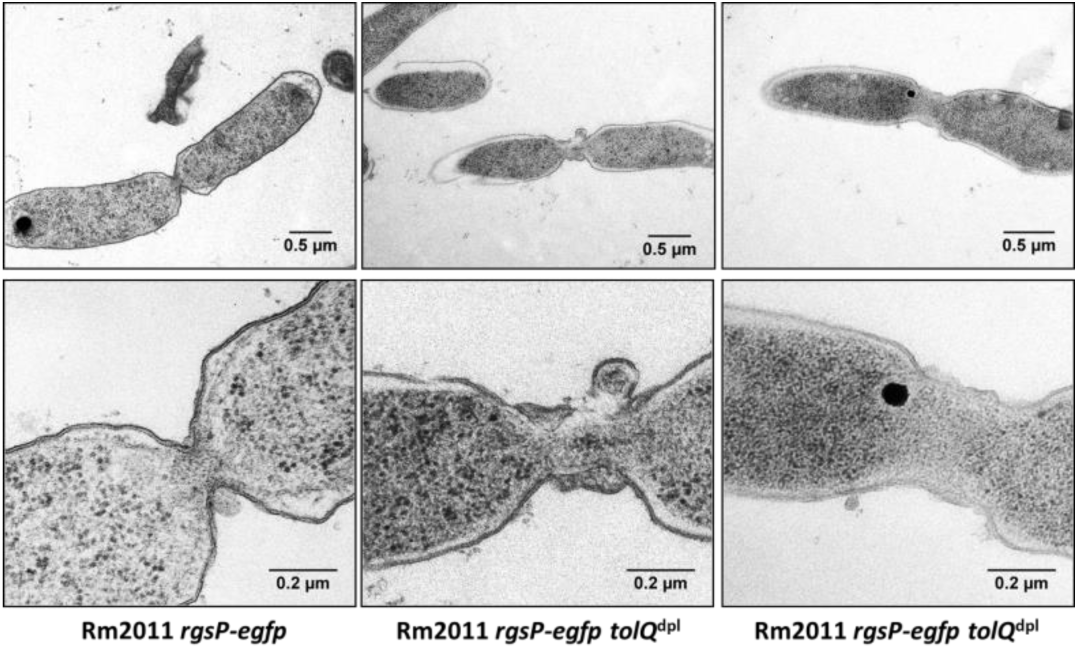
Transmission electron microscopy images of Rm2011 *rgsP-egfp* and its *tolQ* depletion derivative grown in TY without added IPTG for 20 hours.

### Novel Rgs proteins and Tol-Pal localize to cell wall growth zones

In order to determine the subcellular localization of the Tol-Pal system and novel Rgs proteins we replaced *pal* and *rgs* genes at their native genomic locations with versions of these genes encoding fusions to fluorescent proteins. The resulting strains grew normally, indicating that the produced protein fusions were functional. In contrast, replacement of *tolQ* with *tolQ-mCherry* resulted in a growth defect. Therefore, a *tolQ-mCherry* fusion was generated as a single transcription unit driven by a copy of the *tolQ* promoter immediately upstream of the *tolQRAB* operon. Microscopy analysis of these strains in the exponential growth phase revealed polar fluorescence foci, which colocalized with RgsP-EGFP or RgsP-mCherry foci. RgsC-, RgsF-, RgsG- and TolQ-mCherry additionally showed diffuse membrane localization (Fig. 6A). Septal foci of fluorescence were detected for RgsB-, RgsC-, RgsD-, RgsF-, TolQ- and Pal-mCherry as well as for mVenus-RgsS (Fig. 6A). Colocalization of the Tol-Pal system and the novel Rgs proteins with RgsP imply that they accumulate in the areas of cell wall biosynthesis.

**FIG 6.**
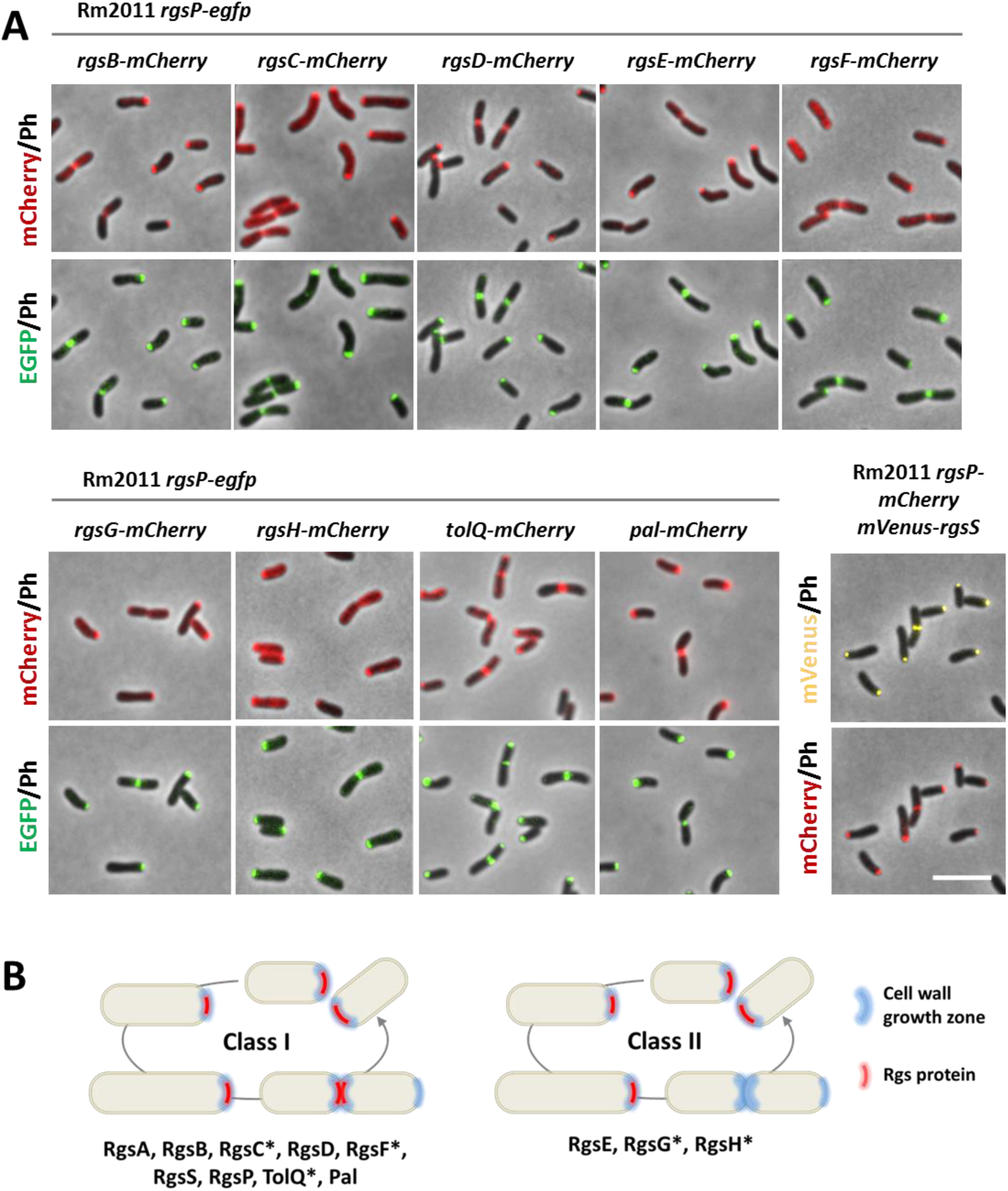
Colocalization of novel Rgs proteins, TolQ and Pal with RgsP-EGFP. (A) Fluorescence microscopy images of Rm2011 *rgsP-egfp* carrying *mCherry* inserted at the native genomic location of *rgsG*, *rgsH*, *tolQ* and *pal* to generate C-terminal mCherry fusions to the respective proteins, and Rm2011 *rgsP-mCherry* carrying *mVenus-rgsS* at the native genomic location, growing exponentially in TY broth. Scale bar, 5 µm. (B) Summary of the time-lapse fluorescence microscopy analysis shown in Fig. S2. Class I Rgs proteins are localized at the growing pole and septum. Class II Rgs proteins are localized at the growing pole only. Asterisk indicates protein fusions that produced diffuse fluorescence signal in addition to localized foci.

The results of the microscopy of exponential culture samples were confirmed in time-lapse studies. RgsB-, RgsC-, RgsD-, RgsF-, TolQ- and Pal-mCherry colocalized with RgsP-EGFP, as well as mVenus-RgsS with RgsP-mCherry over the whole cell cycle. In contrast, RgsE-, RgsG- and RgsH- mCherry were only detected at the growing pole and not at the septum (Fig. S2).

Taken together, these observations suggest that Rgs proteins and the Tol-Pal system are spatiotemporally coordinated. We grouped the Rgs proteins according to their localization pattern. Class I Rgs proteins and Tol-Pal proteins are associated with sites of zonal cell wall biosynthesis during cell elongation and cell division, whereas Class II Rgs proteins are only localized at the site of polar cell elongation (Fig 6B).

### Domain structure and conservation of novel Rgs proteins

Since all novel Rgs proteins were annotated as hypothetical, we applied computational prediction tools to search for conserved domains and characteristic structural features providing first hints at their possible functions (Fig. 2B). Furthermore, we overexpressed the corresponding genes in Rm2011 *rgsP-egfp* and analyzed growth and cell morphology. For this analysis, we used complex TY and LB media. TY medium, containing 2.5 mM CaCl2, is routinely used for cultivation of *S. meliloti* whereas LB medium does not contain CaCl_2_ and was observed previously to potentiate the effect of cell surface-related defects on *S. meliloti* cell morphology (12). Homology and structural similarity analyses of RgsA, RgsB, and RgsH did not provide any hint at their possible functions, whereas RgsG contained conserved domain IalB (COG5342, invasion-associated locus B) (Dataset S2, S3). Overexpression of *rgsA* in Rm2011 *rgsP-egfp* strongly inhibited growth in both media, and caused delocalization of RgsP-EGFP and loss of the rod-shape (Fig. 7). In LB broth, *rgsA*-overexpressing cells were strongly enlarged. Overexpression of *rgsB* in TY-cultivated cells had no effect on cell growth and morphology, as well as on localization of RgsP-EGFP. In cells cultured in LB, *rgsB* overexpression resulted in a phenotype similar to that caused by *rgsA* overexpression. Enhanced expression of *rgsG* and *rgsH* had no effect in either media (Fig. 7).

**FIG 7.**
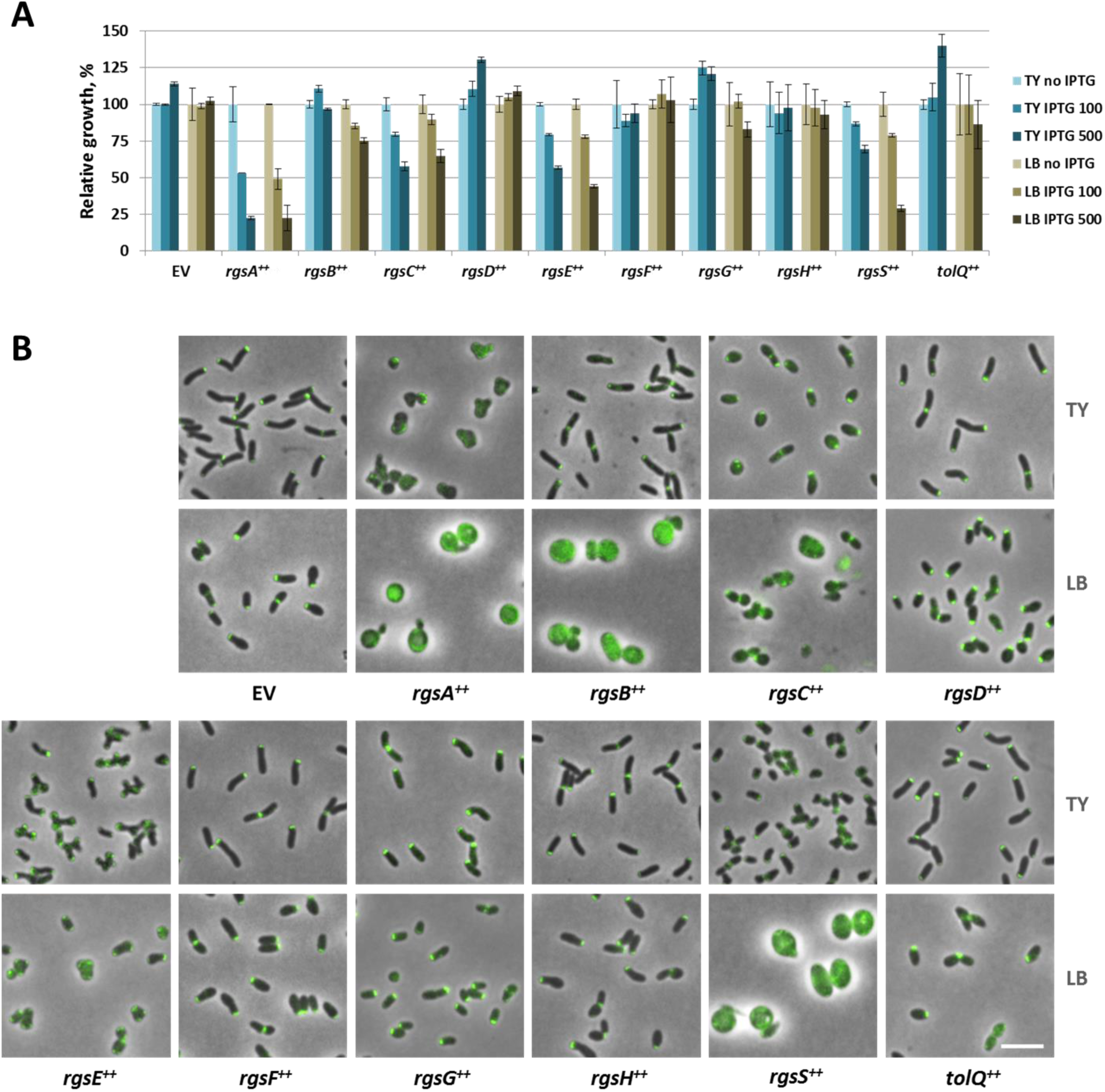
Effects of *rgs* and *tolQ* gene overexpression on cell morphology and growth. (A) Relative growth of Rm2011 *rgsP-egfp* carrying the empty vector pWBT (EV) or *rgs* and *tolQ* overexpression constructs. Precultures grown in TY medium without IPTG were used to inoculate TY or LB cultures with IPTG added at 0, 100 or 500 µm IPTG. The optical density was measured after 16 hours of growth. Values normalized to the growth of the cultures without added IPTG are shown. Error bars indicate standard deviation of three biological replicates. (B) Fluorescence microscopy images of Rm2011 *rgsP-egfp* cells, carrying the empty vector pWBT (EV) or the indicated gene overexpression constructs (*gene*^++^), grown in TY or LB broth with added 500 µm IPTG for 20 to 24 hours. Merged phase contrast and EGFP fluorescence images are shown. Bar, 5 µm.

RgsC contains a conserved M15 Zn metallopeptidase domain in its N-terminal portion (Dataset S2). Structure modelling of RgsC using the SWISS-MODEL online modelling tool (21) identified structural similarities to D-D and L-D carboxypeptidases from Gram-positive bacteria and L-Ala-D-Glu endopeptidase domain of bacteriophage endolysin within the M15 peptidase domain (Dataset S3). Overexpression of *rgsC* in TY-grown cells moderately affected growth and resulted in coccoid cells partially retaining proper RgsP-EGFP localization. In LB broth, the *rgsC* overexpression phenotype included cell lysis (Fig. 7).

RgsE is a 2089 amino acid protein with up to ten Apolipoprotein A1/A4/E domains, homology to SMC chromosome segregation protein (coiled-coil regions) and 42 % of sequence identity to *A. tumefaciens* GPR (Atu1348) (16). Overexpression of *rgsE* in cells cultured in TY broth resulted in cell branching, similar to the morphology of *A. tumefaciens* GPR-overproducing cells. At the cell poles of the branches, polar foci of RgsP-EGFP mediated fluorescence were present. LB-grown *rgsE*-overexpressing cells were slightly enlarged and showed partially delocalized RgsP-EGFP signal (Fig. 7).

RgsF contains a conserved tetratricopeptide TPR domain and six Sel1-like repeats (Dataset S2), suggesting possible functions in protein-protein interactions and signal transduction. Overexpression of *rgsF* did not affect cell growth and morphology in either medium (Fig. 7). The C-terminal portion of RgsS (residues 864-945) constitutes a conserved SPOR domain showing almost equal degree of sequence similarity to the SPOR domains of *E. coli* FtsN and *Bacillus subtilis* cell wall amidase CwlC, whose crystal structures have been reported (22, 23) (Fig. S3A, S3B, Dataset S2). Structure modeling revealed a characteristic arrangement of α-helices and β-strands in the SPOR domain of RgsS (Fig. S3C, Dataset S3). The remaining part of the RgsS amino acid sequence did not deliver significant homology hits pointing to its function. Overexpression of *rgsS* in cells cultured in TY broth resulted in slightly attenuated growth, shorter rods and faint polar RgsP-EGFP foci, whereas in LB medium, growth was strongly reduced and cells were strongly enlarged, displaying a diffuse RgsP-EGFP signal (Fig. 7).

Conservation of the Rgs proteins in α-proteobacterial proteomes was analyzed by an iterative Hidden Markov Model (HMM)-assisted homology search. This revealed that RgsB, RgsC, RgsD, RgsE and RgsF are well-conserved in Rhizobiales, whereas RgsA, RgsH and RgsS were only detected in α-proteobacterial families phylogenetically close to Rhizobiaceae (Fig. 8, Dataset S5). We then employed a manual BLASTP search and identified shorter homologs, sharing similarities with RgsA or RgsS in their C-terminal parts, constituting only a small portion of the cytoplasmic domain, transmembrane helix and periplasmic domain (RgsA) or periplasmic domain (RgsS) (Fig. 8, Dataset S4). Full-length RgsH homologs were ultimately found in 13 out of 16 considered Rhizobiales families (Fig. 8, Dataset S4). In α-proteobacterial orders other than the Rhizobiales, only RgsE and RgsF homologs were found. This analysis indicates that Rgs proteins are almost exclusively conserved in the Rhizobiales, which correlates with the absence of MreB in this order.

**FIG 8.**
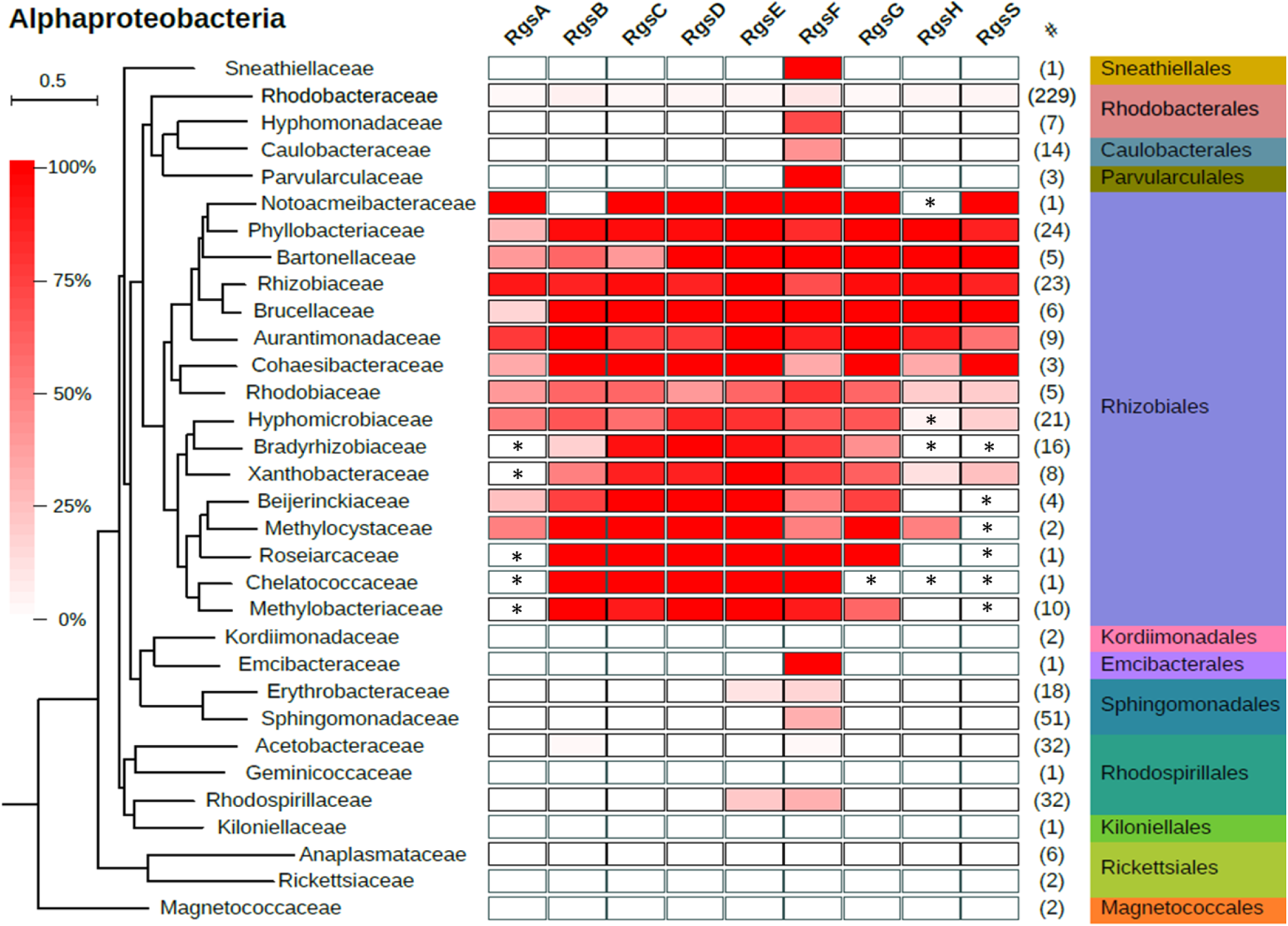
Conservation of novel Rgs proteins in Alphaproteobacteria. The color code indicates the proportion of possessing a respective Rgs protein homolog, grouped at the family level. The number of considered complete proteomes in these groups is shown in column #. The phylogenetic tree was reconstructed using a maximum likelihood approach based on 121 single-copy orthologs common in all considered proteomes. The phylogenetic distance scale is shown at the top left. Asterisks indicate presence of at least one partial homolog, identified in a manual BLASTP search, in a given family.

### RgsD is a novel PG binding protein

A homology search for RgsD did not identify any conserved domains, however modelling the protein structure with SWISS-MODEL revealed structural similarities to isopeptide domains of thioester domain protein BaTIE from *Bacillus anthracis* and ancillary pilin RrgC from *Streptococcus pneumoniae* (Fig. 9A, Dataset S3). These are surface-associated proteins in Gram-positives and suggested to participate in covalent adhesion. Whereas the target of BaTIE binding remained unknown (24), RrgC is covalently attached to the outer PG layer of the streptococcal cell wall (25). We reasoned that RgsD was likely to have affinity to PG and therefore tested whether it interacted with PG. Purified RgsD was retained in the pellet (P) sample following incubation with PG, in a co-precipitation PG binding assay (Fig. 9B). Thus, RgsD represents a novel PG-binding protein.

**FIG 9.**
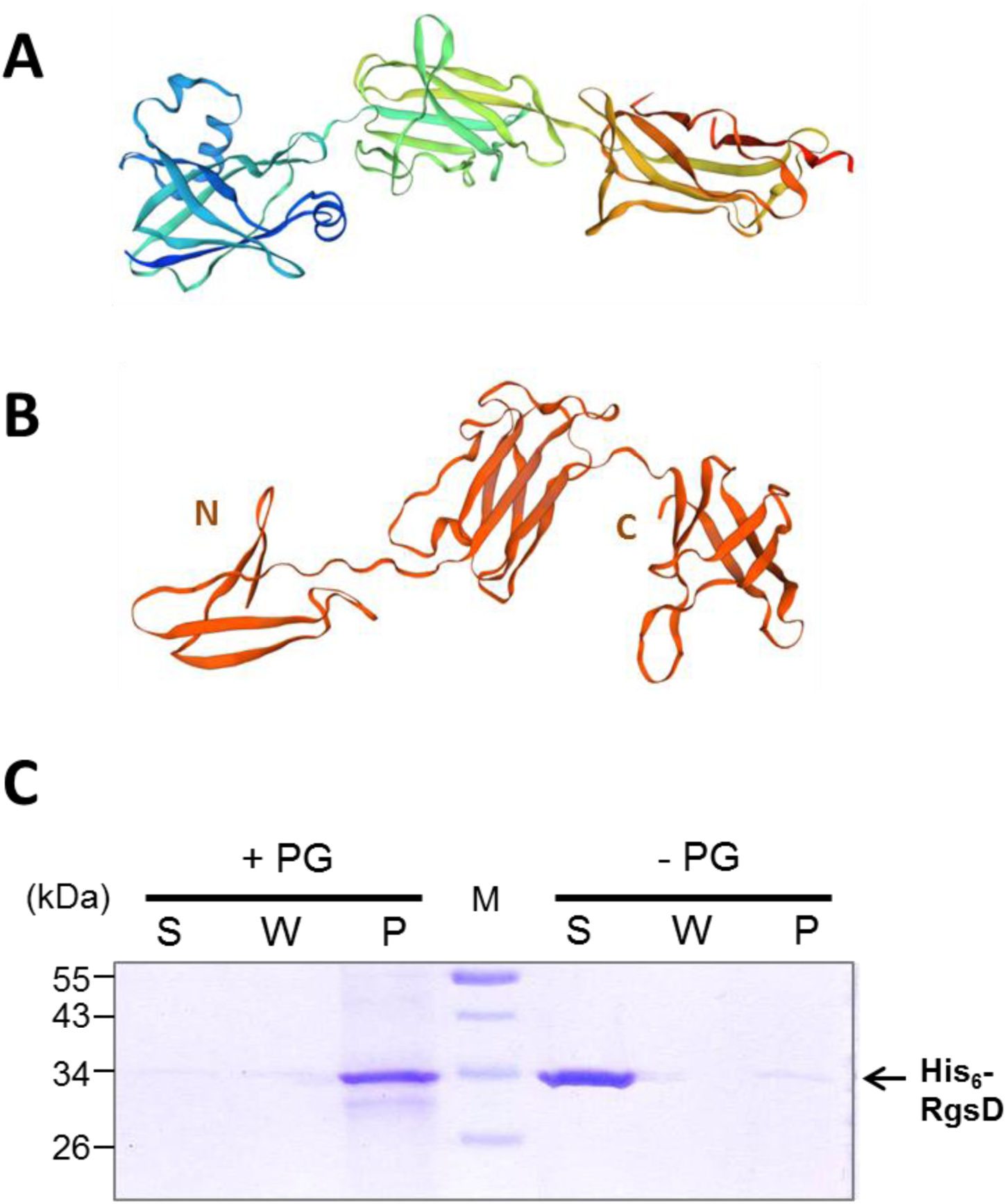
RgsD shows structural similarity to minor pilin RrgC and PG binding ability. (A) Ribbon diagram of experimentally determined RrgC structure (25). (B) Ribbon diagram of RgsD structure model determined by the SWISS-MODEL online tool with RrgC as a template. (C) PG binding ability of His_6_-RgsD_28-331_ was assessed in an *in vitro* binding assay with *S. meliloti* Rm2011 PG sacculi followed by SDS-PAGE and Coomassie blue staining. S, supernatant from the first centrifugation step. W, supernatant from the washing step. P, pellet. PG, peptidoglycan. Control reactions were performed in the absence of PG sacculi.

## DISCUSSION

We previously identified RgsP and RgsM as novel proteins with unknown functions indispensable for *S. meliloti* growth and normal PG composition (12). Correlating with PG incorporation zones, these proteins were detected at the elongating cell pole and relocated to the septum prior to cell division. Here we identified the Tol-Pal system and further six proteins with yet-unknown functions, RgsA, RgsB, RgsC, RgsD, RgsF and RgsS, which followed the spatiotemporal pattern of RgsP and RgsM and were required for *S. meliloti* growth. These represent a set of interacting proteins, associated with zones of cell wall synthesis during cell elongation and cell division, which we define as class I Rgs proteins. In contrast, RgsE, RgsG and RgsH were only detected at the growing pole and not at the septum. These class II Rgs proteins might be specific to the polar cell elongation process.

### Class I Rgs proteins

The periplasmic RgsD protein is exclusively conserved in Rhizobiales and was enriched in RgsA, RgsB, RgsP and RgsM pulldown samples. We identified RgsD as a novel PG-binding protein/domain. PG binding domains are often found in PG-processing enzymes (26). However, the RgsD amino acid sequence did not suggest any enzyme activity. Thus, RgsD could play a role in the positioning of PG synthesis factors or regulation of their activity.

Computational analysis of RgsA and RgsB sequences did not provide any hints to their functions. The sets of Rgs proteins enriched in RgsA and RgsB pulldown samples largely overlapped, suggesting that these two proteins might directly interact with each other. Depletion and overproduction phenotypes of RgsA were more pronounced than those of RgsB. Thus, RgsA possibly has a more important role for cell wall growth than RgsB, which might have an accessory function. Homologs of RgsB were found in all Rhizobiales families except for Chelatococcaceae, whereas RgsA appeared to be less conserved. Full-length RgsA homologs were mostly detected in families phylogenetically close to the Rhizobiaceae. In the remaining Rhizobiales families, shorter proteins with homology to the periplasmic portion of RgsA were present. We therefore speculate that the periplasmic RgsA portion might have a common role in all Rhizobiales families. In the RgsA and RgsB pulldown samples, TolQR and all the Rgs proteins considered in this work were enriched. This implies that RgsAB might be a core component in a Rgs-TolQR protein complex required for *S. meliloti* cell growth and division. This protein complex is likely dynamic, differing in composition between the polar cell elongation zone and the septum.

RgsC, present in all analyzed Rhizobiales families, contains a conserved M15 zinc-binding metallopeptidase domain and shows structural similarities to a DD-carboxypeptidase from Gram-positive *Streptomyces albus* G (27). DD-carboxypeptidases remove the terminal D-alanine residue from penta-muropeptide in PG, thus regulating the abundance of substrates for transpeptidation by the major bifunctional PG synthases (28). In *E. coli*, eight DD-carboxypeptidases contribute to maintenance of the cell shape (29). *S. meliloti* possesses four annotated genes encoding putative DD-carboxypeptidases (*dac*, *SMc00996*, *SMc00683* and *SMc00068*). These contain serine peptidase catalytic domains, characteristic for DD-carboxypeptidases in Gram-negative bacteria. Overexpression of *rgsC* resulted in rounded cells that were prone to lysis in medium without CaCl_2_, indicative of a cell envelope defect. Likewise, overexpression of *dacA* or *dacB* in *E. coli* increased cell envelope permeability (30). It remains to be determined if RgsC has PG carboxypeptidase activity. RgsF represented the only Rgs protein that was substantially conserved outside Rhizobiales. It contains one tetratricopeptide (TRP) domain and six Sel1-like repeats. Domains of this kind mediate protein-protein interactions in all three kingdoms of life (31, 32). *S. meliloti* Sel1-like repeat protein ExoR regulates the essential ExoS-ChvI two-component regulatory system by binding to ExoS (33). Sel1-like repeat proteins were associated with virulence in pathogenic bacteria (34, 35). In *Brucella abortus*, periplasmic protein TtpA containing TRP repeats was required for cell envelope integrity and virulence (36, 37). Interestingly, TPR repeats are present in *E. coli* LpoA, CpoB and NlpI (38–40). LpoA is an outer membrane lipoprotein required for transpeptidase activity of PBP1A, one of the two major PG synthases in *E. coli* (41), the periplasmic protein CpoB regulates the LpoB-mediated transpeptidase stimulation of PBP1B dependent on the Tol energy state (39) and NlpI serves as adaptor protein for PG endopeptidases (40). RgsF does not share sequence homology with *E. coli* LpoA, CpoB or NlpI, however it is a promising candidate for a novel PG synthase activator in polarly growing Rhizobiales.

Out of the six novel class I Rgs proteins, RgsS stood out since its depletion resulted in cell filamentation, suggesting cell division arrest despite ongoing cell elongation. Similar to RgsA, full-length RgsS was only conserved in families phylogenetically close to Rhizobiaceae. In the remaining Rhizobiales families, shorter RgsS homologs with similarities to the C-terminal part of RgsS containing a minor portion of the cytoplasmic domain, transmembrane helix and periplasmic domain, were found. RgsS is a transmembrane protein with conserved SPOR domain at the C-terminus. These features are characteristic of FtsN from γ-proteobacterium *E. coli* and FtsN-like proteins from α, β and δ-proteobacteria, which are highly variable at the sequence level (22, 42). Lesions in FtsN or FtsN-like proteins resulted in cell filamentation due to cell division arrest 42, 43). Thus, RgsS likely represents an FtsN-like protein. In *E. coli*, FtsN interacts with FtsZ-associated protein FtsA and PG biosynthesis-related proteins PBP1B, PBP3, FtsQ and FtsW (44–46). Although *S. meliloti* possesses homologs of these proteins, they were not detected as enriched in the RgsS pulldown assay. In contrast to known FtsN-like proteins, whose cytoplasmic region is short (30 amino acids in FtsN-like CC2007 from *C. crescentus*), RgsS possesses a cytoplasmic domain of 600 amino acids. This suggests an additional functionality related to this cytoplasmic part. One such possible function might be related to cytoplasmic RgsS-RgsE interaction found in our pulldown and bacterial two-hybrid assays.

### Class II Rgs proteins

RgsE is a homolog of *A. tumefaciens* GPR, essential for pole formation in this bacterium (16). RgsE homologs were found in all Rhizobiales families and more distant homologs were detected in Emcibacterales and Rhodospirillales. However, no other Rgs protein homologs were detected in members of these orders. This raises the possibility of a different role for the RgsE homologs in spatiotemporal organization of these bacteria, RgsE and GPR are extraordinary large membrane-anchored cytoplasmic proteins (2089 and 2115 amino acids, respectively), localized to the growing cell pole but absent from the septum. Alterations in cell morphology upon depletion or overproduction of RgsE or GPR in their respective hosts are very similar, supporting a common function of these proteins. RgsE interactions with RgsA, RgsS and TolQ, observed in this work, provide first insights into the mode of RgsE/GRP action in pole formation and polar growth. With accumulating knowledge on the genetics of polar cell wall growth it became apparent that Rhizobiales might employ cell division factors FtsZ, FtsA and FtsW in the restrictive control of polar cell elongation by yet-unknown mechanisms (14, 15). Here, we show that in *S. meliloti* the depletion of another cell division protein, the FtsN-like protein RgsS, resulted in cell filamentation and branching. Relocation of RgsS from the growing pole to the septum might constitute a regulatory cue to inhibit the function of RgsE, ultimately resulting in cessation of polar growth.

Apart from RgsE, the class II Rgs proteins included periplasmic proteins RgsG and RgsH. RgsG is well-conserved in Rhizobiales and contains an IalB domain. This domain was named after invasion-associated locus B from *Bartonella* species, which is an inner membrane protein, promoting the intracellular erythrocyte infection by this pathogen by an unknown mechanism (47). However, RgsG is not a direct homolog of IalB (BARBAKC583_0326) in *Bartonella bacilliformis* KC583, but a homolog of another similar protein, BARBAKC583_0800. RgsH was only partially conserved in Rhizobiales and contains a DUF2059 domain with unknown function. The molecular functions of RgsG and RgsH responsible for their essentiality and localization to the growing pole remain unknown.

### Role of the Tol-Pal system

Identification of TolQ and TolR among the putative interaction partners of RgsA, RgsB and RgsP provided a link between Rgs proteins and known divisome components. Pulldown with TolQ as a bait enriched RgsA, RgsS and RgsE, implying that these proteins might be closely associated with TolQ. The Tol-Pal system is dispensable in *E. coli*, however it was reported to facilitate the cell division process via diverse mechanisms, like maintaining cell envelope integrity, promoting the outer membrane constriction, regulation of PBP1B transpeptidase activity, or indirect activation of amidases acting in septum splitting during cell division (39, 48–50). In the α-proteobacterium *Caulobacter crescentus*, possessing MreB and performing dispersed cell elongation along the sidewall, the Tol-Pal complex is essential. It plays a role for maintaining the outer membrane integrity as well as for successful cell separation (51). Depletion of *S. meliloti* TolQ or Pal resulted in accumulation of normally shaped predivisional cell doublets, suggesting that the final step in cell division was inhibited, but cell elongation and septum constriction proceeded normally. This phenotype was strikingly similar to TolA or Pal depleted *C. crescentus*, which showed doublets of fully developed normally shaped cells (51). Depletion of RgsP and RgsM also resulted in accumulation of constricted pre-divisional cells, which indicated impaired cell division (12). It is tempting to speculate that RgsP, RgsM and possibly further class I Rgs proteins might regulate Tol-Pal-associated processes at the final stages of *S. meliloti* cell division.

We show here that *S. meliloti* TolQ, Pal, and FtsN-like protein RgsS are localized at the growing cell pole and the septum. In *C. crescentus,* Tol-Pal proteins and FtsN-like CC2007 localized at the division plane and were retained at the new cell pole after cell division (42, 51). Absence of cell shape defects in the constricted doublets of TolQ- or Pal depleted cells in both species suggests that these proteins are not required for cell elongation. Likewise, in both species, FtsN-like proteins are unlikely required for cell elongation, considering cell filamentation in their absence. In *C. crescentus* and *A. tumefaciens,* FtsZ-EGFP was detected at the new cell pole (15, 52). It is possible that in some bacteria, such as α-proteobacteria, persistence of cell division proteins at the new cell pole after cell division might be a common feature, which is not related to the cell elongation mode.

## Conclusion

Our survey and initial characterization of RgsP and RgsM interaction partners identified several novel components involved in unipolar cell growth and division in Rhizobiales, and revealed links of these proteins to known cell wall growth and cell division factors. The newly identified Rgs proteins await a detailed analysis of their molecular functions and their interplay to unravel how the complex machineries of the elongasome and divisome accomplish cell growth and division in unipolarly growing α-proteobacteria.

## MATERIAL AND METHODS

### Bacterial strains and growth conditions

Bacterial strains and plasmids used in this study are shown in Table S5. *S. meliloti* was grown at 30 °C in tryptone-yeast extract (TY) medium (53), LB medium (54) or modified MOPS-buffered minimal medium (55). When required, antibiotics were added to agar media at the following concentrations: streptomycin, 600 mg/L; kanamycin, 200 mg/L; gentamicin, 30 mg/L; spectinomycin, 200 mg/L. Unless otherwise indicated, IPTG was added to 500 µM. *E. coli* was grown on LB at 37 °C and antibiotics were added at the following concentrations: kanamycin, 50 mg/L; gentamicin, 8 mg/L; spectinomycin, 100 mg/L. For liquid cultures, antibiotic concentrations were reduced to the half. IPTG was added to 100 µM.

For growth assays involving protein depletion, bacteria were grown in 96-well plates in 100 µl of the medium with or without added IPTG in three replicates, in the Tecan M200PRO instrument, with alternating shaking (10 min) and non-shaking (20 min). Optical density was measured every 30 min. The precultures grown in TY medium with IPTG were used to inoculate the first cultures (Culture I) with or without added IPTG to OD_600_=0.005. After 16 hours of growth, these cultures were diluted in the same media to OD_600_=0.005 to start the second cultures (Culture II) and growth curves were recorded for additional 40 hours.

For growth assays involving gene overexpression, 3 independent transconjugant colonies were used to inoculate 100 µl of TY without IPTG in 96-well plates, followed by incubation with shaking at 1,200 rpm overnight. 1 µl of these precultures were used to inoculate 100 µl of TY or LB medium with or without IPTG as indicated, followed by incubation with shaking at 1,200 rpm for 16 hours. Optical density was measured with the Tecan M200PRO instrument.

For fluorescence microscopy of liquid culture samples and for transmission electron microscopy the *S. meliloti* strains were grown in 3 ml medium in glass tubes, with shaking. Exponential growth phase samples were harvested at OD_600_ between 0.4 and 0.8. For microscopy of protein depletion cultures, precultures in TY medium with added IPTG were used to inoculate TY cultures without added IPTG to OD_600_=0.005 and grown for 20-24 hours. For microscopy of gene overexpression strains, the precultures in TY medium without added IPTG were used to inoculate TY or LB medium with added IPTG (OD600=0.005 if the growth was not inhibited upon gene overexpression and OD_600_=0.01 to 0.05 if the growth was inhibited upon gene overexpression) and grown for 16-20 hours.

### Media

TY medium (5 g/l tryptone, 3 g/l yeast extract, 0.4 g CaCl_2_×2H_2_O). LB medium (10 g/l tryptone, 5 g/l yeast extract, 5 g/l NaCl). MOPS-buffered minimal medium (MM) (10 g/l MOPS, 10 g/l mannitol, 3.55 g/l sodium glutamate, 0.246 g/l MgSO_4_×7H_2_O, 0.25 mM CaCl_2_, 2 mM K_2_HPO_4_, 10 mg/l FeCl_3_×6H_2_O, 1 mg/l biotin, 3 mg/l H_3_BO_3_, 2.23 mg/l MnSO_4_×H_2_O, 0.287 mg/l ZnSO_4_×7H_2_O, 0.125 mg/l CuSO_4_×5H_2_O, 0.065 mg/l CoCl_2_×6H_2_O, 0.12 mg/l NaMoO_4_×2H_2_O, pH 7.2).

### Construction of strains and plasmids

Cloning was performed using PCR, restriction digestion, ligation and *E. coli* transformation. The strains and plasmids generated are listed in Table S1. Primers used in this study are shown in Table S2.

To generate C-terminal fusions to mCherry or FLAG tag sequences encoded at the native genomic location, the C-terminal portion encoding sequence in the range of 500 to 800 bp was inserted in frame into the non-replicative vectors pK18mob2-mCherry or pG18mob-CF and the resulting plasmids were introduced by conjugation into *S. meliloti* strains. This resulted in positioning of the tagged gene copy under the control of the native promoter in the chromosome. To generate the strain with a markerless *mVenus-rgsS* fusion, the mVenus coding sequence was flanked by the upstream non-coding region and an N-terminal portion encoding sequence of the target gene. This construct was inserted into the sucrose selection plasmid pK18mobsacB. Double recombinants were selected on agar medium plates with sucrose (56).

To generate bacterial two-hybrid constructs, the corresponding coding sequences were inserted into the vectors pKT25, pUT18C, pKNT25Spe and pUT18Spe in frame with T25 or T18 fragment encoding sequences of *Bordetella pertussis* adenylate cyclase. Plasmid pKNT25Spe is a derivative of pKNT25, carrying a *Spe*I restriction site directly upstream of the protein-of-interest-T25 fusion translation start, which allows for preserving the native N-terminus of the protein of interest. To construct pKNT25Spe was generated by PCR amplification of pKNT25 with primers shown in Table S2, restriction digestion with *Spe*I and self-ligation.

To generate protein depletion strains, three different strategies were applied. To generate RgsD and RgsF depletion strains, the N-terminal portion encoding sequences, including 27 bp of the upstream non-coding sequence, were cloned into pK18mob2 in the orientation that resulted in placement of the *lac* promoter, carried by the vector, upstream of the partial *rgs* gene. Integration of these constructs into the genome by homologous recombination resulted in placement of the target gene under the control of the *lac* promoter. Subsequent introduction of pWBT, carrying *lacI*, resulted in IPTG-dependent expression of the target gene.

To generate RgsG, RgsH, TolQ and Pal depletion strains, the corresponding gene was deleted from the genome using a pK18mobsac-based deletion construct and the sucrose selection procedure in presence of an ectopic copy of the coding sequence of the gene of interest controlled by the *lac* promoter on plasmid pSRKGm, carrying *lacI*. This resulted in IPTG-dependent expression of the gene of interest.

To generate RgsA, RgsB, RgsC, RgsE and RgsS depletion strains, the corresponding gene was deleted from the genome using a pK18mobsac-based deletion construct and the sucrose selection procedure in presence of an ectopic gene copy including the native promoter on the single-copy curable plasmid pGCH14. This plasmid contains the replication operon *repABC* with a *lacI* box in the promoter region. Therefore, replication of this plasmid can be repressed by LacI. Subsequent introduction of pSRKKm, carrying *lacI*, resulted in IPTG-dependent replication of the complementation plasmid.

Gene overexpression plasmids were constructed by inserting the protein coding sequence into pWBT under the control of the IPTG inducible T5 promoter.

The RgsD protein expression construct was generated using plasmid pWH844 containing sequences encoding an N-terminal His_6_ tag and RgsD lacking the predicted signal peptide.

### Fluorescence microscopy

Microscopy was performed using the Nikon microscope Eclipse Ti-E equipped with a differential interference contrast (DIC) CFI Apochromat TIRF oil objective (100x; numerical aperture of 1.49) and a phase-contrast Plan Apo l oil objective (100x; numerical aperture, 1.45) with the AHF HC filter sets F36-513 DAPI (excitation band pass [ex bp] 387/11 nm, beam splitter [bs] 409 nm, and emission [em] bp 447/60 nm), F36-504 mCherry (ex bp 562/40 nm, bs 593 nm, and em bp 624/40 nm), F36-525 EGFP (ex bp 472/30 nm, bs 495 nm, and em bp 520/35 nm) and F36-528 YFP (ex bp 500/24 nm, bs 520 nm, and em bp 542/27 nm). Images were acquired with an Andor iXon3 885 electron-multiplying charge-coupled device (EMCCD) camera.

For microscopy of exponentially growing cultures, 2 µl of TY cultures at OD_600_ of 0.4 to 0.8 were spotted onto 1 % Molecular biology grade agarose (Eurogentec) pads, let dry for 2-3 minutes, closed with cover glass and microscoped. For time-lapse microscopy, bacteria from exponential growth phase TY cultures were diluted 1:20 and 2 µl were spread by gravity flow on the MM agarose pad and let dry for 14 minutes. The pads were closed air-tight with the cover slip and microscoped in an incubation chamber at 30 °C.

Cell morphology analysis was performed using the MicrobeJ plugin to the ImageJ software. Graphical representation of the data and statistics analysis was performed with the PRISM software (GraphPad) using t-test with Welch correction.

### Transmission electron microscopy

Sample preparation and microscopy were performed as described previously (12). Briefly, concentrated *S. meliloti* cell suspensions were high pressure frozen and impregnated with substitute resin. The polymerized resin blocks containing the samples were cut to 50 nm ultrathin sections using an ultramicrotome and applied onto 100 mesh copper grids coated with pioloform. The sections were imaged using a JEM-2100 transmission electron microscope (JEOL, Tokyo, Japan) equipped with a 2k x 2kF214 fast-scan CCD camera (TVIPS, Gauting, Germany).

### Protein purification

His_6_-RgsD was purified as described previously (57). Briefly, cells were lysed using M110-L microfluidizer (Microfluidics) and the protein was isolated using 1-mL HisTrap column (GE Healthcare). Protein was concentrated with Amicon Ultracel-10K (Millipore) to a volume of 2 ml and applied to size exclusion chromatography (SEC; HiLoad 26/600 Superdex 200 pg, GE Healthcare), After SEC, protein containing fractions were pooled and concentrated with an Amicon Ultracel-10K (Millipore) according to the experimental requirements.

### Co-immunoprecipitation (Co-IP) and protein identification by mass-spectrometry

Co-IP and protein identification by mass spectrometry was performed as previously described including small modifications (58). Briefly, *S. meliloti* cultures, producing the C-terminal FLAG fusions and the negative control strain Rm2011 carrying the empty vector pWBT were grown in 200 ml of TY medium to an OD_600_ of 0.4-0.6 cross-linked with 0.36% formaldehyde for 15 min at RT and then quenched for 10 min with glycine (final concentration of 0.35 M). Cells were lysed using French press instrument. Cleared lysates were obtained after centrifugation for 1 hour at 20,000 x g and incubated with anti-FLAG M2 affinity gel (FLAG Immunoprecipitation Kit, Sigma). Bound proteins were eluted with 3xFLAG peptide solution.

Mass-spectrometry analysis to identify eluted proteins was performed as described previously (12). The data were filtered to exclude proteins identified with less than two unique peptides. The total signal was calculated as a sum of area values. Relative protein enrichment was calculated as a percent area value of the total signal multiplied by coverage value. The data from experimental samples was further filtered to cut off proteins with the relative enrichment value less than 0.5.

### Bacterial two-hybrid analysis

Bacterial two-hybrid analysis was performed as previously described (59). The adenylate cyclase-deficient strain *E*. *coli* BTH101 was co-transformed with corresponding plasmid pairs. Single cotransformant colonies were inoculated in 100 μl LB supplemented with ampicillin and kanamycin and grown at 30 °C for 6 hours with shaking at 1,200 rpm. 10 μl of each culture was spotted onto LB agar plates containing kanamycin, ampicillin, 40 mg/L X-Gal and 500 µM IPTG. Plates were imaged after 20 hours of incubation at 30 °C and 24 hours of incubation at room temperature. 3 independent cotransformant colonies were analyzed and produced similar results.

### Peptidoglycan purification and muropeptide analysis

*S. meliloti* strains were grown in TY with IPTG overnight to obtain the precultures. These were used to inoculate 400 ml TY without IPTG to OD_600_ of 0.001 (2011 *rgsP-egfp*) or 0.005 (2011 rgsP-*egfp rgsA*dpl) and grown for 20 hours at 30 °C. Isolation of PG sacculi was performed as described previously (12).

Purified sacculi (∼10 µg) were digested with the muramidase Cellosyl and the resulting muropeptides were analyzed by HPLC as previously described (12). The muropeptides were assigned according to established nomenclature (60).

### Peptidoglycan binding assay

Purified PG (∼100 μg) from *S*. *meliloti* strain Rm2011 was centrifuged at 15,000 ×*g*, 4 °C for 14 min and resuspended in binding buffer (10 mM Tris-maleate, 10 mM MgCl_2_, 50 mM NaCl, pH 6.8). 10 μg of protein of interest was incubated with or without PG in a final volume of 100 μl, and incubated at 4 °C for 30 min. Samples were centrifuged as described above and the supernatant was collected (supernatant fraction) whilst the pellet was resuspended in 200 μl of binding buffer. Another centrifugation step as described above was carried out (wash fraction). Bound proteins were released from PG by incubation with 100 μl of 2% SDS, at 4 °C for 1 hour before being collected by a final centrifugation step as done at earlier steps. The proteins present in the different fractions were analyzed by SDS-PAGE.

### Homology search

To assess the distribution of Rgs proteins in bacteria, 3,835 proteomes of the UniProt “Reference Proteomes 2019-05” set were searched (61). This subset does not include species with no or preliminary taxonomic classification (unclassified, candidatus, sp.) and species that represent subspecies of otherwise present species (genosp., subsp.). The Rgs homologs from *S. meliloti* 2011, *Agrobacterium tumefaciens* (strain C58 / ATCC 33970), *Mesorhizobium loti* MAFF303099 and *Rhodopseudomonas palustris* TIE-1 (exception: no RgsH and RgsS known in this species) were used as references.

For each Rgs protein an iterative search was performed based on profile hidden Markov models of the reference sequences using *jackhmmer* (62). This initial search was limited to Alphaprotebacteria (552 species) and run with an E-value threshold of 1e-40 and five iterations. Results were filtered automatically in two stages. The first stage was the size of the amino acid sequence which should be within +/-25% of the reference protein of *S. meliloti* 2011 (an exception is the large and repeat-containing RgsE protein for which a threshold of +/-50% was used). The second stage was the annotation of transmembrane topology and signal peptides using the predictor tool Phobius (19). Assuming that the majority of putative homologs is correct, sequences with a transmembrane topology deviating from the most frequent one of the respective Rgs protein were removed. The remaining protein sequences were aligned progressively using MUSCLE (63) to create a largely improved profile hidden Markov model. This model was then used in a single search run against proteomes of all bacterial classes using HMMsearch (64). MUSCLE-based alignments of all predicted homologs are shown in Dataset S5.

### Phylogeny of Alphaprotebacterial families

For each family of Alphaprotebacteria one representative species was selected by choosing the species with the smallest taxonomy ID in the UniProt Reference Proteomes. *E. coli* (TaxID: 83333) was used in addition as an outgroup. The phylogenetic reconstruction was performed as describer previously (65). The amino acid sequences of all 121 single copy orthologs universal to this set identified by Proteinortho (66) with an e-value of 1e^-40^ were aligned using MUSCLE and concatenated to a 57,845 aa long “super protein”. Before this C and N-termini of the aligned sequences were trimmed until no gaps were left. ProtTest3 (67) identified the LG matrix with a Gamma model of rate heterogeneity, an estimate of proportion of invariable sites and empirical base frequencies (+I+G+F) (68) as the best model according a Bayesian Information Criterion (BIC). Hence, a rapid bootstrap analysis with 1000 replicates was performed to search for best-scoring maximum likelihood tree with respect to this model using RaxML with *E. coli* as outgroup (69).

### Data availability

All co-immunoprecipitation data are presented in Supplemental dataset S1. Features from computational analysis of Rgs proteins are presented in Supplemental datasets S2 and S3. Sequence comparisons underlying conservation analysis of Rgs proteins in Alphaproteobacteria are shown in Supplemental datasets S4 and S5.

## ACKNOWLEDGMENTS

We thank Dr. Uwe Linne for support of MS-based protein identification and Dr. Thomas Heimerl for preparation of samples for transmission electron microscopy.

